# Differential effect of climate on structural and biochemical plant traits

**DOI:** 10.1101/2022.07.18.500466

**Authors:** Dinesh Thakur, Věroslava Hadincová, Renáta Schnablová, Helena Synková, Daniel Haisel, Nada Wilhelmová, Tomáš Dostálek, Zuzana Münzbergová

## Abstract

Exploring patterns and causes of intraspecific trait variation is crucial for a better understanding of the effects of climate change on plant populations and ecosystems. However, our current understanding of the intraspecific trait variation is mainly based on structural (morphological) traits, and we have limited knowledge on patterns and causes of variation in biochemical traits (e.g., leaf pigments), which are also crucial for plant adaptation. As a result, we also do not know how similar the climatic effects on structural versus biochemical traits are.

Using plant traits from 110 genotypes representing 11 *Festuca rubra* populations grown in 4 different climates, we studied trait covariation among structural traits (linked to fitness, resource use, gas exchange, and reproduction) and biochemical traits (linked to photosynthesis, photoprotection, and oxidative stress). We also disentangled the relative role of the climate of origin and the climate of cultivation in the structural versus biochemical traits and tested for adaptive plasticity in the traits.

We found that 1) biochemical traits did not covary with structural traits and represent independent *‘photoharvesting – photoprotection’* strategy dimension of functional variation; 2) interactive effects of climate of origin and cultivation were more pronounced for biochemical than structural traits. 3) Trait plasticity was affected by the climate of origin (precipitation and temperature as well as their interaction); 4) *F. rubra* showed both adaptive and mal-adaptive plasticity, and adaptiveness depended upon trait type, cultivation climate, and climate of origin.

Overall, our results suggest that structural and biochemical plant traits respond differentially to climate and thus the response of one group of traits cannot be predicted from the other. Responses are also strongly determined by interactions between the climate of origin and cultivation. Thus, more studies on variation in biochemical traits, their correspondence to other traits, and their variation with climate are needed.

## INTRODUCTION

Plant traits determine various plant functions, define ecological strategies and describe plant responses to environmental change and their role in ecosystem functioning (Westoby *et al*. 2002; Violle *et al*. 2007; Enquist *et al*. 2015; Kattge *et al*. 2020). Therefore, establishing links between functional traits and environmental factors is crucial for understanding how plant populations and communities respond to climate change.

Traits vary both within and between species. However, our trait-based understanding is primarily based on interspecific trait variability, and only recent studies focused on the implications and consequences of intraspecific trait variation (Bolnick *et al*. 2011; Siefert *et al*. 2015; Westerband *et al*. 2021). Lately, intraspecific trait variation has been suggested to be equally important as interspecific variation for understanding, predicting, and managing large ecological processes (Siefert *et al*. 2015; Henn *et al*. 2018; Westerband *et al*. 2021).

Intraspecific trait variation arises from both genetic differences and phenotypic plasticity (Matesanz *et al*. 2012). Intraspecific variation in traits due to genetic differences is achieved very slowly by natural selection, while plasticity allows for rapid responses. Many studies have elucidated genetic versus plastic responses in multiple plant traits in various species (Münzbergová *et al*. 2017; Datta *et al*. 2017; Kosová *et al*. 2022). However, most of these studies deal only with structural traits (i.e., morphological traits, e.g., specific leaf area, plant height, plant biomass). Variation in biochemical traits, such as leaf pigment content, enzymes, and primary and secondary metabolites, is an important mechanism through which plant individuals can rapidly increase their fitness in a variable environment (Taiz *et al*. 2015), but variation in such traits is much less studied. Unlike structural traits, biochemical traits have low costs, are reversible and also faster to produce. Thus, biochemical traits could be more plastic in response to environmental variability than structural traits. For example, in response to temperature variability over a season, various biochemicals such as leaf pigments and sugars accumulate several times during cold acclimatization, revert back during de-acclimation and are reused for growth during favourable conditions (Liu *et al*. 2019; Rathore *et al*. 2021).

Biochemical traits are closely related to physiological activity of plants under variable environments (Taiz et al. 2015) and should be closely related to structural traits, as all structures in plants are formed as a result of biochemical processes (e.g., high content of photosynthetic pigments should lead to increased plant biomass). The available ecological studies exploring biochemical traits study them separately of structural traits (see, e.g., Alonso-Amelot et al. 2007, Martin et al. 2007). Due to this, we do not know whether the genetic versus plastic responses differ between structural and biochemical traits. Identifying specific plant functions that are determined more by genetic variation or are more plastic is crucial for our ability to understand and predict species responses to the ongoing climate change more accurately (Henn *et al*. 2018; Pérez-Ramos *et al*. 2019).

Phenotypic plasticity is considered a major mechanism of response to environmental variability (Valladares *et al*. 2006; Nicotra *et al*. 2010; Henn *et al*. 2018). Plasticity is advantageous (i.e., adaptive) only if it increases fitness of an individual in a new environment (van Kleunen and Fisher 2001; Nicotra *et al*. 2010). Despite an increasing number of studies on the adaptiveness of plasticity (e.g., in Palacio-López et al. 2015, Acasuso-Rivero et al. 2019, Brooker et al. 2022), most of the studies are considering only a few plant traits indicative of plant fitness (e.g., seeds produced, clonal individuals or number of flowers) and rarely incorporate biochemical plant traits (but see Arnold *et al*. 2022). Estimates of adaptiveness of plasticity is problematic in perennial plants with frequent vegetative reproduction, where fitness estimates are not straightforward. Recently, Enquist et al. (2015) suggested that we can determine whether phenotypic plasticity is adaptive by exploring if the trait changes toward its optimum in a specific environment (i.e., trait values in the home environment). If the traits of plants grown in a foreign climate converge towards traits of plants which are at home in that climate, the trait shift is considered adaptive. However, only very few studies have attempted to explore this so far. For example, Henn *et al*. (2018) provided evidence that transplanting plant species to a new community result in an adaptive shift in leaf trait values. Similarly, Challis *et al*. (2022) reported adaptive plasticity in traits related to hydraulic failure in a tree species.

The extent and adaptive nature of trait plasticity likely depends on the conditions of plant origin as well as on conditions in which they are currently growing (Valladares *et al*. 2006). For instance, plants from one climate of origin may be more plastic and plastic responses may be stronger to one environmental factor than to another factor. Münzbergová *et al*. (2017) showed that plants originating from drier sites show more plasticity in structural traits. However, we do not know how this effect of climate of origin on plasticity differs among structural and biochemical traits. Considering our expectation related to higher genetic differentiation in structural traits, it can be expected that plasticity of structural traits will be affected more by climate of origin than biochemical traits.

Because plasticity is known to depend upon climate of origin as well as on climate of cultivation (Münzbergová *et al*. 2017; Kreyling *et al*. 2019), plants from some climate of origin grown in a given environment may show adaptive plasticity while plants from other climate may show mal-adaptive plasticity in the same conditions (Ghalambor *et al*. 2007). For example, Price et al. (2003) suggested that when the climate to which an individual is exposed differs markedly from climate of origin, the plasticity is more likely to be mal-adaptive. However, such knowledge is still largely lacking. Therefore, we need to elucidate if the adaptive or mal-adaptive plasticity is consistent among populations from different climates of origin shifted to different novel climates. We also need to test if adaptive plasticity is consistent among different trait types (e.g., structural versus biochemical traits). This will enable us to better understand the adaptive potential of plant populations under climate change.

At both intraspecific and interspecific levels, traits do not vary independently but covary with other traits (Westoby *et al*. 2002; Wright *et al*. 2004; Yang *et al*. 2019). The covariation among traits is due to multiple traits having common functional roles or developmental pathways, or due to genetic linkages and pleiotropy (Armbruster and Schwaegerle 1996). Examining the trait covariation improves understanding of plant ecological strategies and fundamental trade-offs (e.g., acquisitive – conservative resource use strategy) and can also help to improve the prediction of one set of traits from others (e.g., root morphology from leaf morphology) (Westoby *et al*. 2002; Yang *et al*. 2019). However, we don’t know about trait covariation patterns (and plant adaptation strategies) at biochemical levels and if structural and biochemical traits covary in the same dimensions.

Considering the above gaps, we used plant genotypes sampled from 11 populations of *Festuca rubra* occurring in contrasting climates (climates of origin) and grown in 4 different growth chambers (cultivation climates) and measured a wide range of biochemical and structural traits of the plants to address the following questions:

1. What are the major dimensions of variation in structural and biochemical traits (trait covariation)?
2. Do the relative role of climate of origin and cultivation climate and their interactions in trait variation differ among structural and biochemical traits?
3. What is the magnitude of variation in structural and biochemical traits in plants from different climate of origin in response to contrasting cultivation climates?
4. Is plasticity in both structural and biochemical traits adaptive and does the adaptiveness depend upon climate of origin and cultivation climate?

We hypothesized that 1) some of the structural traits covary with biochemical traits (e.g., plant biomass and content of photosynthetic pigments), because plant structural traits are the result of processes occurring at biochemical levels; 2) biochemical traits are more plastic (and less genetically controlled) and thus less affected by the climate of origin than structural traits because of their flexible nature and ease in construction; 3) both structural and biochemical traits show adaptive plasticity, but biochemical traits show a higher degree of adaptive plasticity. This is again because of their lower construction costs and flexibility that can be altered within short timescales.

## METHODOLOGY

### Study species and sampled localities for plant material

In the experiment, we used *Festuca rubra* ssp. *rubra*, a widespread perennial grass species that occurs in temperate grasslands in Europe. This species can reproduce both by seeds and clonally by rhizomes. The experimental plants of this species were collected in 2014 from 12 natural localities representing factorially crossed precipitation and temperature gradients corresponding to the SeedClim Grid (Meineri *et al*. 2014) in Western Norway. This grid represents an independent combination of three levels of mean summer temperature (4 warmest months, 6.5 °C, 8.5 °C and 10.5 °C) with four levels of annual precipitation (600, 1300, 2000 and 2700 mm).

The plant material used in the current study has been collected for previous studies, that have set up the experiments and provided data also for this study (Münzbergová et al. 2017). From each locality, at least 40 plants of *F. rubra* (at least 1 m apart) were collected in the growing season of 2014 and transported to the experimental garden of the Institute of Botany, Czech Academy of Sciences in Průhonice, Czech Republic (49°59’38.972”N, 14°33’57.637”E) and planted into pots. To ensure that each of the collected plants corresponded to a single genotype, the plants were reduced to a single ramet and planted into 16 × 16 × 16 cm pots in 1:2 soil:sand ratio in August 2014. Later, the plants were checked by flow cytometry and confirmed that all belonged to the hexaploid cytotype of *F. rubra* ssp. *rubra* (Šurinová *et al*. 2019). In one of the localities (i.e., 6.5 °C temperature and 1300 mm precipitation), *F. rubra* ssp. *rubra* was absent. So, finally, we worked with plants from only 11 localities.

In November 2014, 25 genotypes from each locality were transferred to a greenhouse with temperature between 5 and 10 °C. In the greenhouse, the genotypes were multiplied (for details see (Münzbergová *et al*. 2017)). At the end of February 2015, four ramets of each of the 25 genotypes from all the 11 localities were planted in separate pots, resulting in a total of 1100 ramets. One ramet of each genotype was shifted to one of four different cultivation climates (see next paragraph) so that each genotype is present in each cultivation climate.

### Cultivation climates

The experimental plants were grown in four different growth chambers (Vötsch 1014) under conditions simulating four extreme localities in the SeedClim grid (wettest/driest combined with warmest/coldest) for the spring to summer climate in the field. The temperature and moisture in each of the growth chambers followed the same course as in the simulated natural locality (for details, see Münzbergová et al., 2017). The moisture level in the growth chambers was monitored using TMS dataloggers (Wild *et al*. 2019) (three inside each chamber) and then adding the necessary amount of water to mimic the soil moisture of the natural localities (also monitored using the TMS dataloggers). Briefly, in the climate chambers with dry conditions, plants were watered with about 20 ml of tap water per plant, applied to the trays if the soil moisture was lower than 15%. In the wet regime, plants were cultivated under full soil saturation with ~1.5 cm water level in the tray. By manipulating the soil moisture in the growth chambers, soil moisture at localities with a certain precipitation level was mimicked. Thus, the moisture conditions in the growth chamber were referred to as precipitation values of the simulated localities. Day length and radiation levels inside the growth chambers were also controlled in a manner to mimic the conditions of the original localities (Münzbergová *et al*. 2017).

Due to the limitation of resources to take measurements of the biochemical traits from all the grown plants (see below), only 10 genotypes per locality were used for the purpose of this study. The other genotypes along with those considered in this study were used in other studies exploring the growth, fitness, and physiological traits of the plants (Münzbergová *et al*. 2017; Stojanova *et al*. 2018; Kosová *et al*. 2022; Thakur and Münzbergová 2022). The dataset used in this study comprised of samples collected from 440 plants (i.e., 11 localities × 10 genotypes × 4 cultivation climates).

### Plant trait measurements

From each genotype in each cultivation climate, we measured 11 biochemical traits and 8 structural (i.e., morphological) traits. The biochemical traits were represented by the content of antheraxanthin, ß-carotene, chlorophyll a, chlorophyll b, phenolic substances, lutein, neoxanthin, superoxide dismutase (SOD), violaxanthin and zeaxanthin and xanthophyll cycle de-epoxidation state (DEPS). These biochemical traits have been estimated specifically for this study. The considered biochemical traits are related to plant photosynthesis, photoprotection and oxidative stress response (Table 1).

**Table 1:** List of structural and biochemical traits used in this study and their significance along with loadings of trait variation explained by the first five principal components (PCs). Traits with # in superscript were cube root transformed and with $ were log transformed prior to analysis. PC loadings for SLA are missing because they were estimated from genotypes from only 4 localities and not used in overall PCA.

| Trait | Role | PC1 | PC2 | PC3 | PC4 | PC5 |
| --- | --- | --- | --- | --- | --- | --- |
| **Structural traits** |  |  |  |  |  |  |
| Aboveground biomass | Allocation to above ground resource acquisition; indicator of plant fitness | -0.34 | **1.40** | -0.01 | 0.72 | 1.18 |
| Plant height (cm)<sup>#</sup> | Plant competitive ability for light | -0.59 | 0.83 | 0.33 | -0.36 | **1.59** |
| Rhizome weight (g)<sup>#</sup> | Investment into clonal reproduction | **-1.02** | 0.10 | -0.88 | -0.07 | -0.32 |
| Root weight (g)<sup>#</sup> | Allocation to soil resource acquisition | **-1.23** | 0.92 | -0.67 | 0.47 | -0.36 |
| Specific leaf area (SLA, mm<sup>2</sup>/mg) | Per unit leaf construction cost; related to plant resource use strategy | - | - | - | - | - |
| Stomatal density (number/ 500 × 500 µm leaf area) | Gas exchange; allow plants to adjust their stomatal pore area in response to the surrounding environment. | -0.15 | **-1.27** | -0.96 | 0.06 | 0.87 |
| Stomatal size (µm) |  | -0.21 | 1.28 | 0.66 | -0.10 | **-1.00** |
| Number of ramets<sup>\$</sup> | Measure of plant fitness and related to plant resource use strategy | -0.32 | 0.51 | **-1.58** | 0.77 | -0.35 |
| **Biochemical traits** |  |  |  |  |  |  |
| Chlorophyll *a* (µg/g) | Key photosynthetic pigments in plants | **1.97** | 0.25 | -0.42 | -0.34 | 0.07 |
| Chlorophyll *b* (µg/g) |  | **1.43** | 0.11 | -0.62 | -0.87 | 0.22 |
| β-carotene (µg/g) | Responsible for the quenching of light and protection of cells from damages caused by light and superoxide radicals | -0.26 | 0.41 | -0.04 | **-1.59** | -0.04 |
| Antheraxanthin (µg/g)<sup>#</sup> | Accessory light-harvesting pigments and plays important role in the protection of photosynthetic machinery | **-1.83** | 0.01 | -0.18 | -0.50 | 0.07 |
| Neoxanthin (µg/g)<sup>#</sup> |  | **1.61** | 0.44 | -0.54 | -0.06 | 0.08 |
| Violaxanthin (µg/g) |  | **1.73** | 0.41 | -0.45 | -0.12 | -0.14 |
| Zeaxanthin(µg/g) |  | **-2.00** | -0.10 | -0.18 | -0.19 | -0.03 |
| Lutein (µg/g) |  | **1.95** | 0.17 | -0.15 | -0.08 | -0.01 |
| Xanthophyll cycle de-epoxidation state (DEPS)<sup>#</sup> | De-epoxidation state provides information on xanthophyll cycle activity | **-2.09** | -0.17 | -0.09 | -0.14 | 0.00 |
| Phenolic compounds (µg/g) | UV protection by scavenging the reactive oxygen species | -0.79 | 0.41 | -0.37 | -0.97 | -0.14 |
| Superoxide dismutase activity (SOD)<sup>\$</sup> | First line of defence against oxidative stresses; role in scavenging the reactive oxygen species | -0.66 | 0.05 | -0.70 | -0.55 | 0.01 |

Structural traits included aboveground biomass, plant height, rhizome weight, root weight, stomatal density, stomatal size, and the number of ramets. Additionally, specific leaf area (SLA) was measured from the 10 genotypes originating from each of the four most extreme climates of the climatic grid. All these structural traits have been also used in our previous studies to address totally different hypotheses (Münzbergová *et al*. 2017; Kosová *et al*. 2022; Thakur and Münzbergová 2022). The considered structural traits are linked to plant resource use, competitive ability, fitness, growth and gas exchange (Table 1). All the traits were estimated following the standard methodology described in Münzbergová et al. (2017), Münzbergová and Haisel (2019), and Kosová et al. (2022) and in Supplementary Methods.

### Data analysis

All statistical analyses were performed using R version 4.0.5 (R Development Core Team 2019). SLA was estimated only for genotypes from 4 localities (corners of SeedClim grid), and therefore it was only used in analyses involving traits from genotypes from only these localities and not used in any of the following analyses involving genotypes from all the localities.

#### Trait covariation analysis and dependence of traits on climate

We used principal component analysis (PCA) on standardized trait values [‘vegan’ package (Oksanen *et al*. 2013)] to characterize trait covariation patterns and to identify the major dimensions of functional variation. The number of principal components to retain after PCA was determined on the basis of randomization-based procedure (Peres-Neto *et al*. 2005) using rndLambdaF function of ‘PCDimension’ package (Wang *et al*. 2018). Our data had 39 missing values (0.5 %). Therefore, prior to analysis, missing values in the data were imputed using iterative principal component analysis (PCA) algorithm by implementing the imputeFAMD() syntax in the ‘missMDA’ package (Josse and Husson 2016). This imputation of missing values was important specifically for doing multivariate analyses. To check the effect of data imputation on the PCA results, we also conducted PCA on the original data after omitting the samples with missing values. As there were only very minor differences among both PCAs (see Supplementary Table 1), only the one based on the imputed dataset is explored in the main text.

To analyze the dependence of traits on temperature and precipitation of original and cultivation climate and their interaction, we used redundancy analysis (RDA, vegan package) and tested their effects using permutation tests. In this analysis, all the variables and their interactions had significant effect on traits, but some had only very low explanatory power. Therefore, in the final model, we only retained the variables which were significant and explained at least 1% variation in the data. A table showing the change in R^2^ upon addition of the given predictor to the model along with significance is shown in Supplementary Table 2. We also explored impacts of the directionality of the climate differences by subtracting the values of temperature and precipitation of the original localities from the values of temperature and precipitation in growth chambers following Münzbergová *et al*. (2017). We then tested the effect of these differences and their interaction on plant traits using RDA.

#### Relative role of different factors in variation of structural versus biochemical traits

To investigate the relative role of each factor (temperature and precipitation of origin and cultivation climate and their interactions) in trait variation, we determined what fraction of the total variance was explained by each factor independently. This was done for each trait individually by performing variance decomposition analysis using linear mixed-effects model in ‘lme4’ package. In these models, the intercept was the only fixed effect and all the factors (i.e., temperature and precipitation of origin and cultivation climate) and their interactions, as well as genotype, were entered as random effects. The variance attributable to each factor was then expressed as a percentage of the total variance. Such models for partitioning the total variance into various components have been widely used (Messier *et al*. 2017; Isaac *et al*. 2017; Anderegg *et al*. 2018; Henn *et al*. 2018). Before analysis, the traits with non-normal distributions were transformed appropriately (log or cube root, see Table 1).

The variance partitioning results obtained from the analyses (Supplementary Data) above were then used to test if the effect of climate of origin, cultivation climate and interactions between the climate of origin and cultivation differ among trait types. The variance due to climate of origin was taken as the sum of variation explained by original temperature, original precipitation and their interaction (Otemp, Oprec and Otemp × Oprec in Supplementary Table 3), due to the cultivation climate as variance explained by cultivation temperature, cultivation precipitation and their interaction (Ctemp, Cprec and Ctemp × Cprec in Supplementary Table 3). The variance due to all the remaining interactions represents the variance due to the interaction of climate of origin and climate of cultivation (indicating genetic differentiation in plasticity). The effect of trait type on explained variance by climate of origin, cultivation climate and their interaction was analysed using the Mann-Whitney test with explained variance as a response variable.

We also estimated the plasticity (measure of magnitude of variation) of each trait for each genotype across all the four cultivation climates as

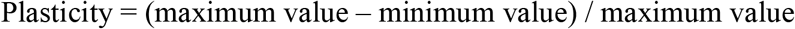

These plasticity values were then used to test the effect of climate of origin on trait plasticity. The relationship between the climate of origin and plasticity of all the traits was analyzed using RDA with plasticity values as the response variable and climate of origin (temperature and precipitation and their interaction) as explanatory variables.

#### Trait adaptive potential

To estimate trait adaptive potential, we performed a PCA analysis (in the same way as in the section ‘trait covariation analysis’), but separately for biochemical and structural traits and expressed the position of each genotype (in each growth chamber separately) as its PC scores. For structural traits, we used scores of the first 3 axes and for biochemical traits, we used scores of the first 2 axes (scores are given in Supplementary Table 4). In structural traits, the first three axes of the PCA explained 30.2%, 19.3% and 17.4% of the total variance, respectively. For biochemical traits, the first two axes explained 55.7% and 15.4% of the total variance, respectively. These scores later served as summarised biochemical and structural traits. We then tested the effects of climate of origin and cultivation climate and their interaction on these scores using linear mixed-effect models with genotype as a random factor in ‘lme4’ package (Bates *et al*. 2015) in R (model statistics in Supplementary Table 5). This test has only been done for the four extreme populations corresponding to the climates simulated in the growth chambers and thus also included SLA.

Based on these models, we calculated marginal means of the summarised traits for plants of each origin in each cultivation climate. We then estimated standardized effect size among pairs of means and tested which pairs of means differed significantly from each other. Both marginal means and standardized effect sizes were estimated using ‘emmeans’ package (Searle *et al*. 2022) in R. We used this information to make inferences if traits converged or diverged or none of the two in response to cultivation climate. For example, let us consider two climates A and B. In case of convergence, when plants originating from A will be grown in B climate their traits will be more similar to plants of origin B grown in B climate than the traits of plants originating in A and grown in A. Divergence means the opposite. Therefore, in case of trait convergence among plants (from different climates of origin) grown in a common climate, there will be a significant shift in the estimated marginal mean of plants grown in a foreign climate towards that of plants that are grown in their climate of origin (decreased standardized effect-size). According to Enquist et al. (2015), phenotypic plasticity may be adaptive if it results in the shift of trait values toward an optimum that is adaptive for a specific environment, i.e., a trait can be considered as adaptative if we detect trait convergence, while trait divergence indicates mal-adaptation.

## RESULTS

### Trait covariation

The PCA showed that the studied traits are not independent but show covariation patterns. Our analysis revealed 5 independent dimensions of trait variation and structural traits varied independently of biochemical traits. The first 5 axes of the PCA accounted for 66% of the total trait variation (Table 1 and Supplementary Figure 1). PC1 alone explained more variation than the other 4 axes combined and was dominated mainly by biochemical traits related to photosynthetic machinery (photosynthetic traits as opposed to photoprotective traits). Root weight also varied along PC1 but orthogonally to biochemical traits (Figure 1). PC2 represented traits related to plant fitness and biomass allocation and gas exchange. Number of ramets had strongest loading along PC3, β-carotene along PC4 and size-related traits along PC5 (details in Table 1 and Supplementary Figure 1).

**Figure 1:**
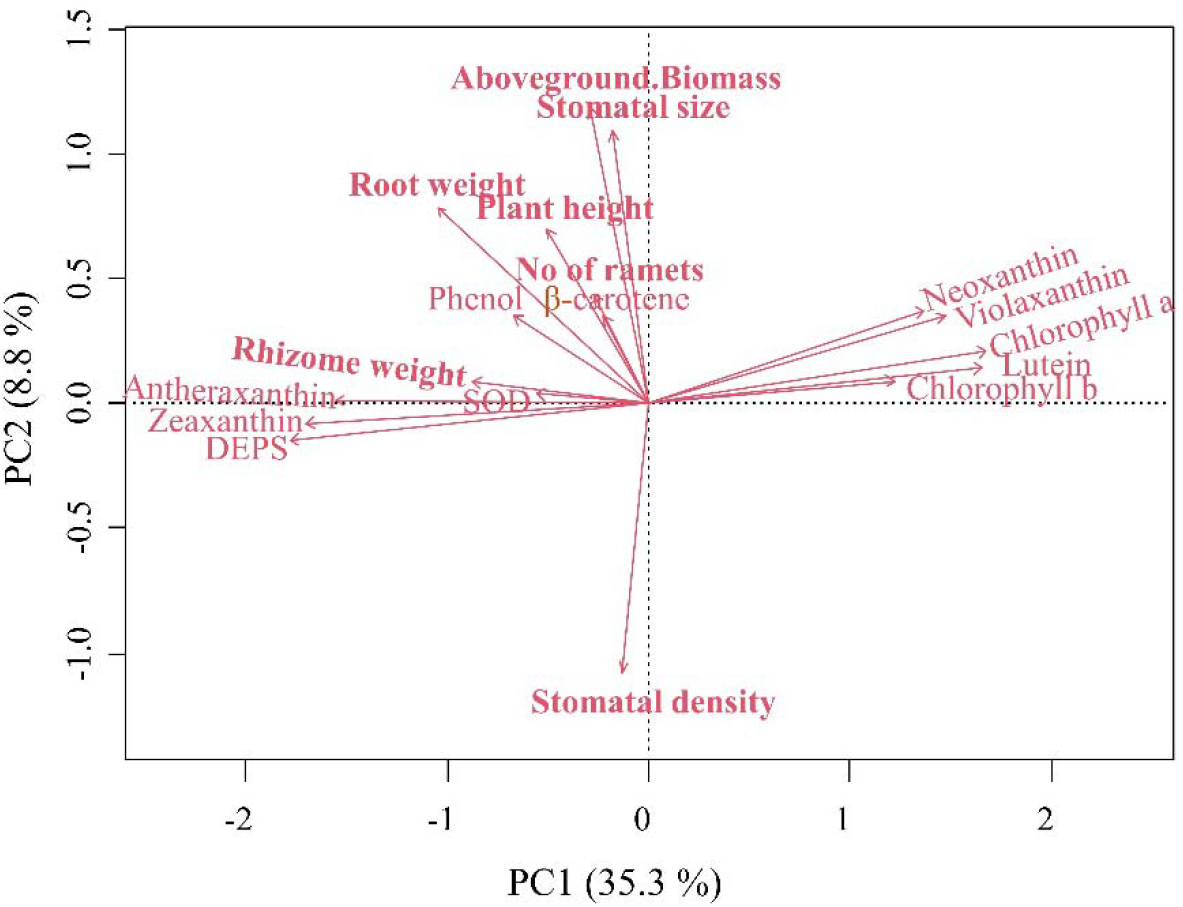
Principal component analysis showing covariation patterns of studied structural (in bold) and biochemical traits. DEPS = xanthophyll cycle de-epoxidation state; SOD = Superoxide dismutase.

### Effect of original and cultivation climate on traits

The RDA analysis showed that the climatic factors have a significant effect on the traits (Supplementary Table 6) and explained 37% of their total variation. The first two axes jointly explained 31 % of the variation, with axis 1 explaining 8 times more than axis 2. The precipitation of the cultivation climate (Cprec) was the key variable along with the interaction of Cprec with the precipitation of origin (Oprec) and the temperature of the cultivation conditions (Ctemp) associated with axis 1 (Figure 2). Ctemp along with Cprec × Ctemp were the variables associated with axis 2. The traits related to photosynthetic machinery increased in the wetter cultivation climate, whereas the traits related to photoprotection and biomass of belowground parts tend to have higher values in plants cultivated in drier climates. The key variables that tend to increase with increasing temperature of the cultivation climate were plant height and aboveground biomass.

**Figure 2:**
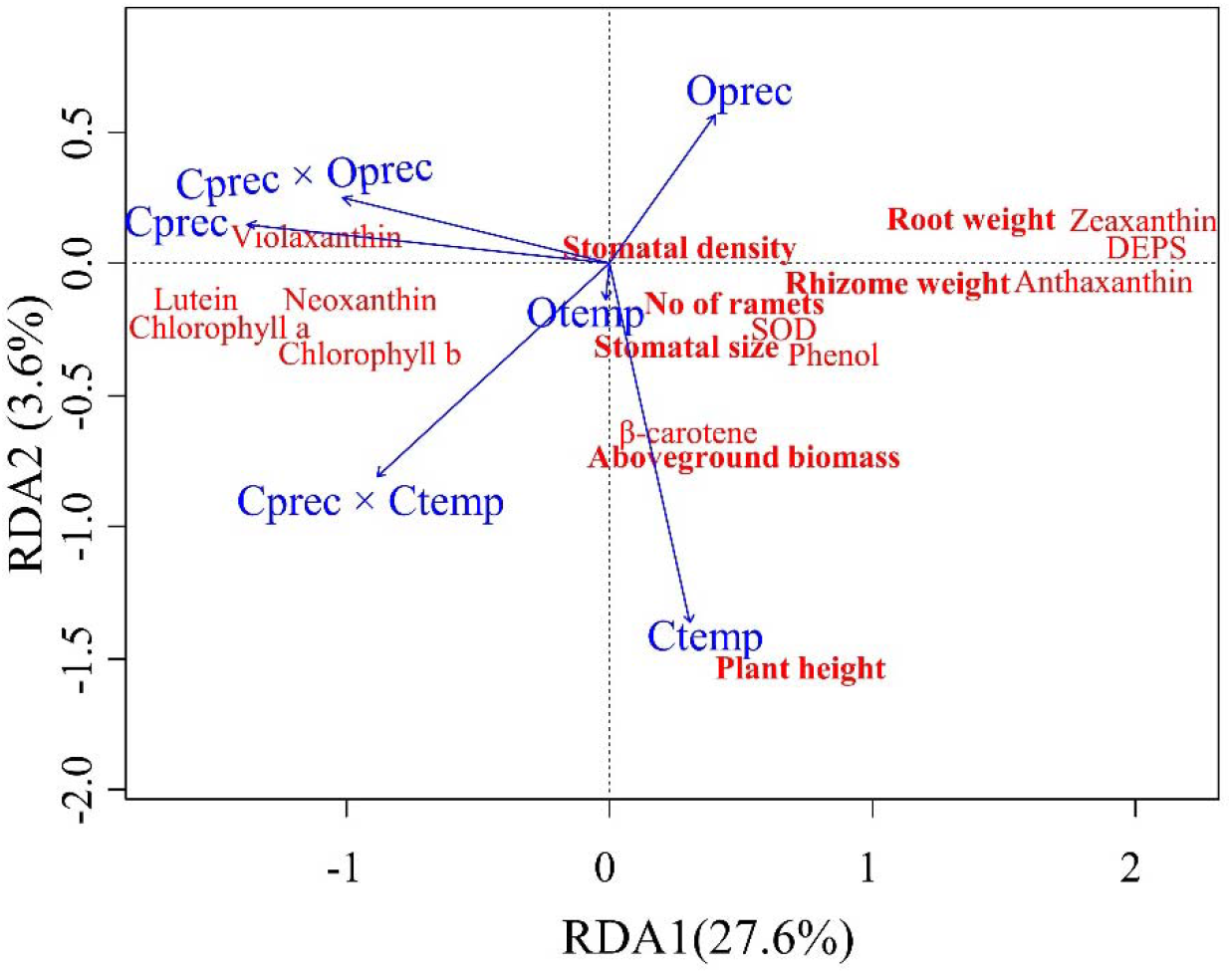
RDA plot representing the effects of cultivation climate and climate of origin on structural (in bold) and biochemical traits. DEPS = xanthophyll cycle de-epoxidation state; SOD = Superoxide dismutase. Oprec and Otemp/Cprec and Ctemp represent precipitation and temperature of climate of origin/cultivation climate.

Climate differences [difference between precipitation and temperature of cultivation and the origin] had a significant effect on traits (Supplementary Table 7). In the RDA with differences in precipitation and temperature as predictors, 23% of the total variance in traits was explained (Figure 3). Axis 1 explained a lot more (10×) variation than axis 2. The difference in precipitation was associated with axis 1 and the difference in temperature was associated with axis 2. The interaction between the difference in precipitation and the difference in temperature was not associated with either of the first two axes. From the RDA plot, it is evident that photoprotective pigments and their allocation to the lower parts (roots and rhizomes) increased while photo-harvesting pigments decreased in plants shifted to drier conditions (Figure 3). Plant height increased in plants shifted to warmer conditions.

**Figure 3:**
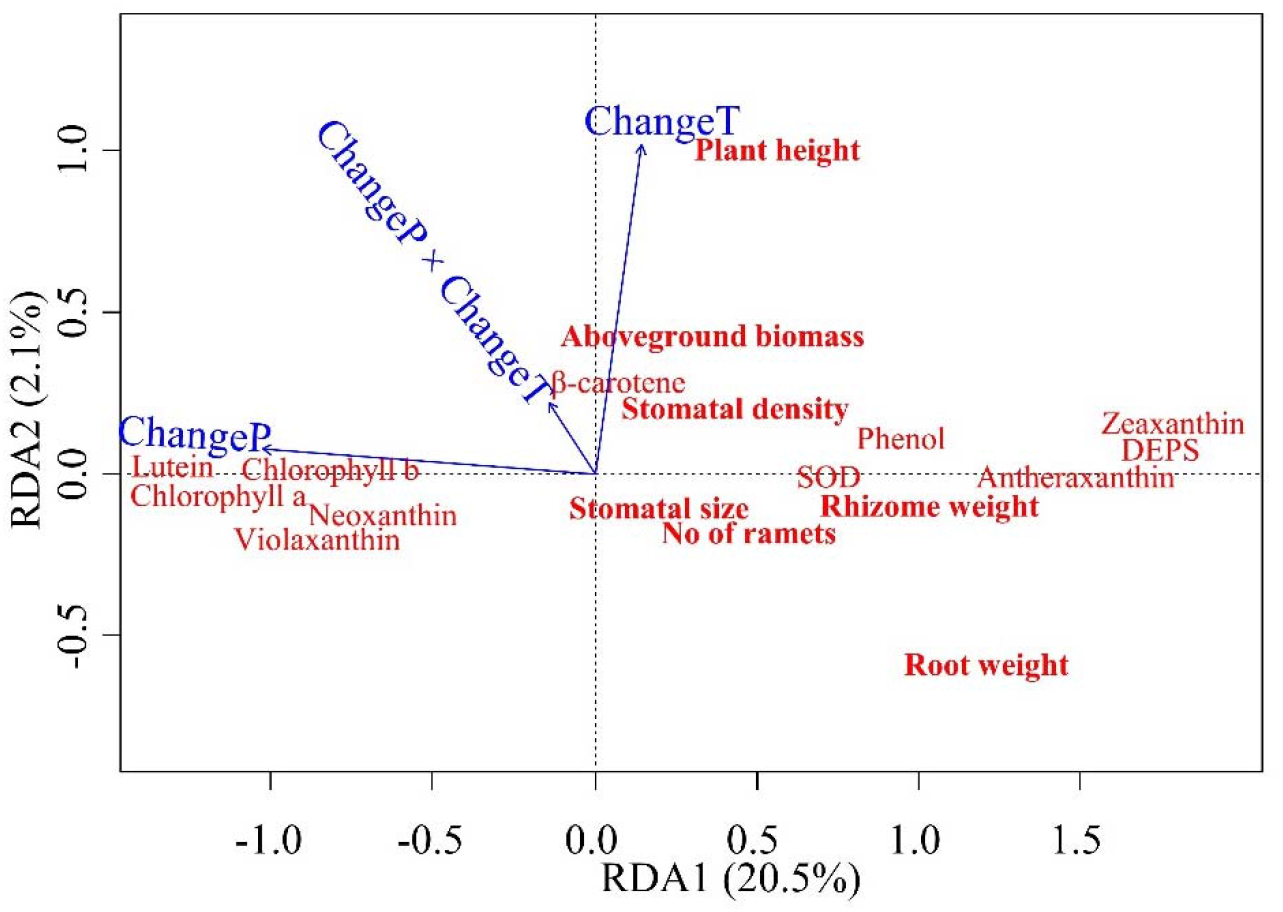
RDA plot representing effect of climate differences (cultivation - origin) on structural (in bold) and biochemical traits. DEPS = xanthophyll cycle de-epoxidation state; SOD = Superoxide dismutase. ChangeP and ChangeT represent difference in precipitation and temperature between origin and cultivation climate (cultivation-origin).

### Drivers of trait variation in structural versus biochemical traits

The relative role of climate of origin and its interactions with the climate of cultivation on traits was specific to trait type (p < 0.05 in the Mann-Whitney test), while the variation due to cultivation climate was not significantly different (p = 0.27). The variation explained by genotype also differed significantly among trait types and variation explained by genotype was more in structural traits (p = 0.05). Variation in structural traits due to the interaction of climate of origin with cultivation climate was significantly lower than in biochemical traits (p < 0.001). The opposite was observed for variation due to the climate of origin (p = 0.049) (Figure 4). The interaction between the climate of origin and cultivation climate indicates that plastic responses depend upon the climate of origin (i.e., genetic differentiation in plasticity). Therefore, our results suggest that there is higher genetic differentiation in plasticity in biochemical traits.

**Figure 4:**
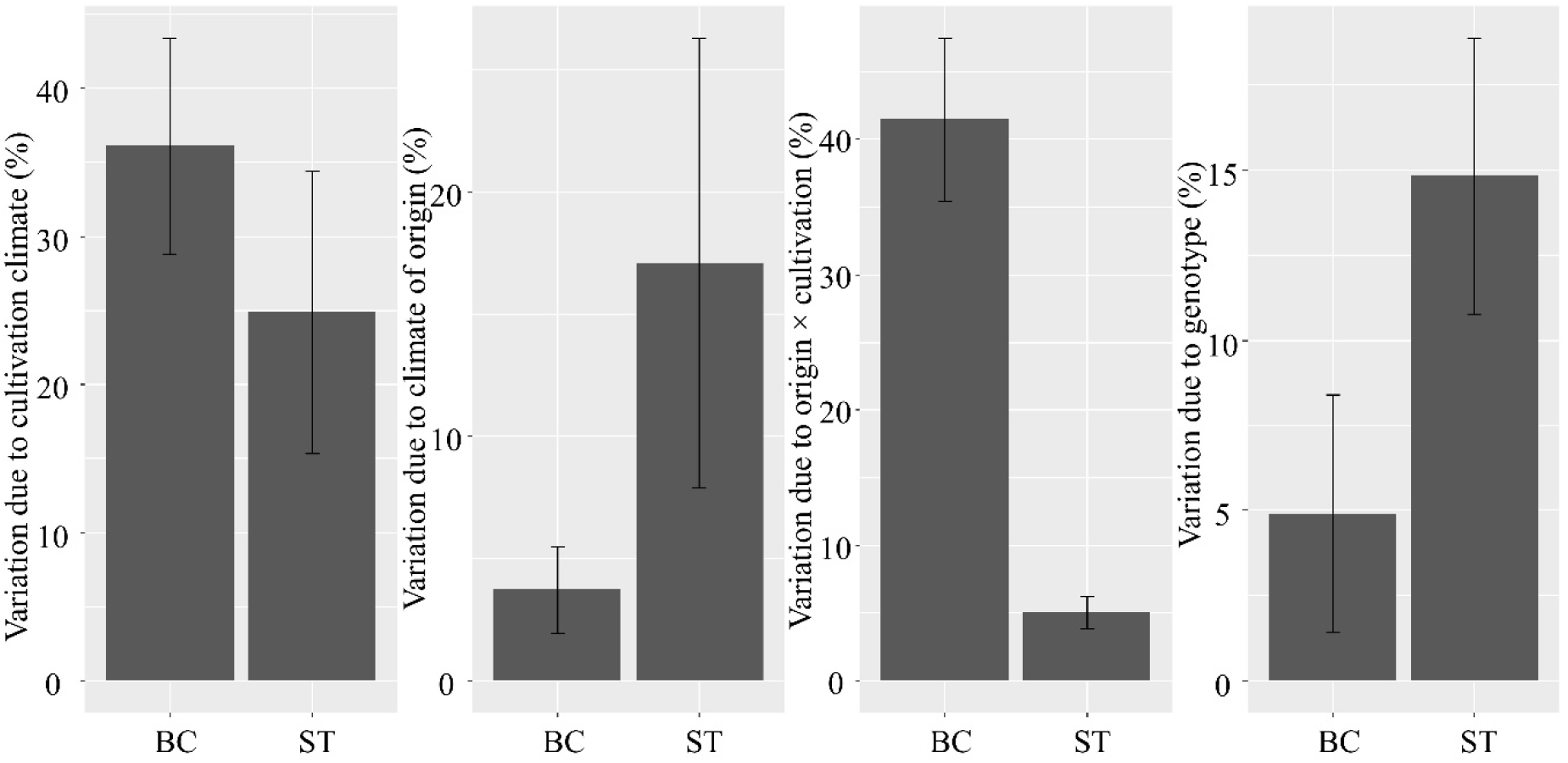
Comparison of the fraction of total variation explained by cultivation climate, climate of origin, their interaction and genotype in biochemical (BC) versus structural (ST) traits. Means and their standard errors are shown. This figure is based on Supplementary Table 3.

Genotypes from different climates of origin differed in their plasticity. In the RDA, a total of 19 % of the variability in the plasticity of traits was explained by the climate of origin. Both precipitation and temperature of origin, as well as their interaction, significantly affected trait plasticity (p < 0.05). The biochemical traits (except phenol and SOD) of plants originating from wet or wet and warm climates tend to be more plastic than the biochemical traits of plants originating from drier climates (Figure 5).

**Figure 5:**
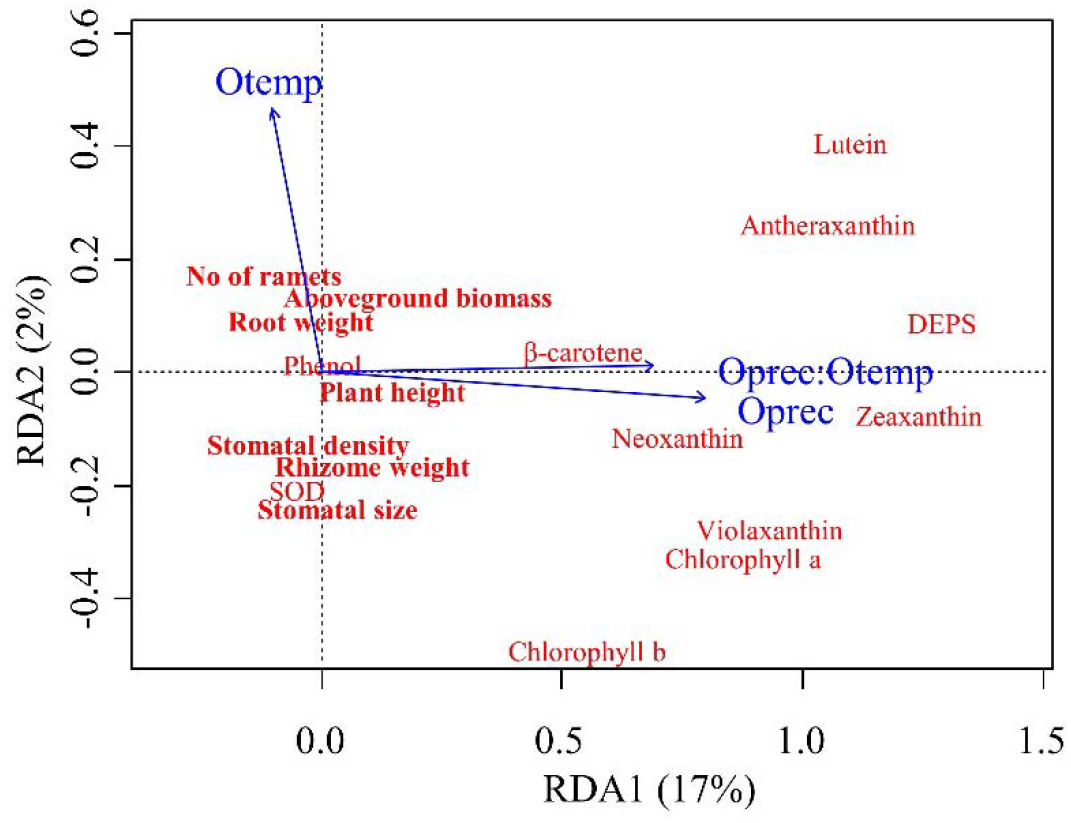
RDA plot representing the effect of climate of plant origin on plasticity in structural (in bold) and biochemical traits. DEPS = xanthophyll cycle de-epoxidation state; SOD = Superoxide dismutase. Oprec and Otemp represent precipitation and temperature of climate of origin.

#### Trait adaptive potential

In structural traits, the general tendency was to converge (i.e., the traits showed adaptive potential). PC1 (based only on structural traits) which was dominated by plant fitness traits (i.e., number of ramets and plant biomass traits, see Supplementary Table 4) showed the highest degree of convergence (Table 2, Supplementary Figure 2) while PC2 scores (representing gas exchange traits) mainly showed no change. The results based on PC3 (representing plant size) showed either convergence or no change. Biochemical traits representing PC1 (photoharvesting and photoprotection related traits) tend to diverge except convergence in cold and wet cultivation climates (Table 2, Supplementary Figure 3) while those representing PC2 (related to low wavelength light harvesting) tend to converge. Overall, these results suggest that *F. rubra* shows adaptive plasticity as well as mal-adaptive plasticity in response to different cultivation climates. However, adaptive plasticity depends upon trait type, cultivation climate and climate of origin.

**Table 2:** Trait convergence or divergence in different populations of *Festuca rubra* when grown in different foreign climates. Inferences were made based upon the significant differences between estimated marginal means between plants in climate of origin and cultivation climate. Convergence refers to significant shift in estimated marginal mean (based on PCA scores generated using traits) of plants grown in a cultivation climate towards that of plants in climate of origin. Conv. = convergence; Div. = divergence; None = neither convergence nor divergence. Cells with ‘-’ represent those plants were in climate of origin. This table is based on mixed effects model statistics reported in Supplementary Table 5 and is based on analysis using genotypes originating only from the four most extreme localities.

|  |  | Cultivation climate |  |  |  |
| --- | --- | --- | --- | --- | --- |
|  |  | Cold and dry | Warm and dry | Cold and wet | Warm and wet |
| Climate of origin |  | Structural traits (PC1) |  |  |  |
|  | Cold and dry | - | None | Conv. | Conv. |
|  | Warm and dry | None | - | Conv. | Conv. |
|  | Cold and wet | Conv. | Conv. | - | Conv. |
|  | Warm and wet | None | Conv. | None | - |
|  |  | Structural traits (PC2) |  |  |  |
|  | Cold and dry | - | None | None | None |
|  | Warm and dry | None | - | None | None |
|  | Cold and wet | None | None | - | None |
|  | Warm and wet | Conv. | None | None | - |
|  |  | Structural traits (PC3) |  |  |  |
|  | Cold and dry | - | None | Conv. | None |
|  | Warm and dry | None | - | None | None |
|  | Cold and wet | Conv. | Conv. | - | Conv. |
|  | Warm and wet | Conv. | None | None | - |
|  |  | Biochemical traits (PC1) |  |  |  |
|  | Cold and dry | - | None | Conv. | Div. |
|  | Warm and dry | Div. | - | Conv. | Div. |
|  | Cold and wet | Div. | Div. | - | None |
|  | Warm and wet | Div. | Div. | Conv. | - |
|  |  | Biochemical traits (PC2) |  |  |  |
|  | Cold and dry | - | None | Conv. | Conv. |
|  | Warm and dry | Conv. | - | Conv. | Conv. |
|  | Cold and wet | Conv. | Conv. | - | Conv. |
|  | Warm and wet | None | Conv. | None | - |

## DISCUSSION

### Independent covariation of biochemical traits and the *photoharvesting – photoprotection* strategy dimension

Contrary to our expectation, the trait covariation analysis revealed independent variation of structural and biochemical traits in the multivariate trait space. The possible reason for the decoupling of the two groups of traits could be distinct strategies and differential effects of climatic at biochemical and structural levels. This lack of covariation and trait variation into multiple independent dimensions limits our ability to predict biochemical plant functions from morphological plant functions or *vice versa*. Therefore, our results highlight the importance of incorporating biochemical plant traits (which are largely ignored) in ecological studies to better understand plant functioning in response to environmental change.

The second key finding emanating from the covariation analysis was the covariation between photosynthesis and photoprotective biochemical traits along the first axis. At the positive end of this axis, there were individuals with high photoprotective and low photosynthetic pigments, a syndrome that can be considered as the *‘photoharvesting – photoprotection’* strategy dimension of functional variation [similarly to the famous *‘conservative-exploitative’* resource use strategy based on the leaf economic spectrum, (Wright et al. 2004)).

The strategy we propose represents a trade-off among leaf biochemical traits related to photosynthetic machinery and is characterized by an increase in photoharvesting pigments with a decrease in photoprotective pigments or *vice versa*. Higher amounts of photoprotective pigments are important in high light conditions for the protection of photosynthetic machinery (Ramírez-Valiente *et al*. 2015; Munné-Bosch *et al*. 2016) and high amounts of photoharvesting pigments are important in low light conditions to make maximum use of available solar radiation. In this study, all genotypes were grown under the same light conditions in each cultivation climate simulating the original light conditions in Norway; hence, light is not the direct driving factor here. The variation in leaf pigments in this study may be determined by temperature and precipitation (of both origin and cultivation climates). For instance, plants growing in higher temperatures and drier conditions have a higher chance of photodamage (Taiz *et al*. 2015; Szymańska *et al*. 2017) and hence higher amounts of photoprotective pigments are needed even if the light is unchanged.

Another trade-off evident in our analysis was between stomatal size and stomatal number, which has been demonstrated also in range of earlier studies (e.g. Lawson and Blatt 2014; Kardiman and Raebild 2018). In this study, stomatal size also positively covaried with plant aboveground biomass. Smaller and greater number of stomata may cause loss of excessive water through transpiration leading to sub-optimal gas exchange and lost opportunities for photosynthesis (Lawson and Blatt 2014), possibly explaining the covariation observed.

### Effects of climate differences on structural and biochemical traits

Temperature is often considered more important than precipitation in climate change context (Bjorkman *et al*. 2018; Vitasse *et al*. 2018). However, our results suggested that the difference in precipitation (cultivation – original climate) had a stronger effect on plant traits (i.e., caused a greater plastic response) than the difference in temperature. The effects were stronger in biochemical traits, which are only rarely studied in climate change context [accept studies of crops or a few experimental species (Hekneby *et al*. 2006; Pradhan *et al*. 2019; Arnold *et al*. 2022)]. Photosynthesis related traits were the most affected by climate differences (cultivation – original climate), suggesting that the most important process leading to primary production in the Biosphere is the most responsive (i.e., shows plastic responses) to climate change. The high effect of the difference in precipitation on photosynthesis related traits could be because photosynthesis is closely related to gas exchange through the stomata and their opening and closing depends upon water availability (Taiz *et al*. 2015; Kardiman and Raebild 2018).

Plant height and aboveground biomass increased, and root mass decreased when plants were shifted to warmer climate (grown in warmer climate than climate of origin). This finding is similar to previously reported in a metanalysis by Lin *et al*. (2010). The reason for increased aboveground biomass under warmer climates could be attributed to increased rate of photosynthesis and decreased thermal restriction of tissue formation (Hoch *et al*. 2002). The possible reason for decreased allocation to root mass when plants were shifted to warmer and wetter climates could be linked to increased resource availability [water and nutrient availability in soil, due to increased mineralization] (Pugnaire *et al*. 2019). The increased root mass was also linked to decrease in precipitation of cultivation. This is the expected result as more roots are needed to meet the water demand in drier climates. Overall, these findings highlight the importance of studying trait plasticity in response to changes in precipitation along with temperature and incorporating biochemical traits in studies to better understand the impacts of climate change on plants.

### Interactive effects of climate of origin and cultivation climate are stronger on biochemical traits

High intraspecific trait variation is considered important in population persistence (Bolnick *et al*. 2011; Siefert *et al*. 2015) and can be achieved through genetic differentiation and/or plasticity (Matesanz *et al*. 2012). We expected structural traits to be more genetically differentiated with higher genetic differentiation in plasticity (i.e., the stronger role of interactions between the climate of origin and cultivation) than biochemical traits. We found that structural traits were more genetically differentiated (stronger effect of climate of origin), but biochemical traits were more strongly affected by the interaction of the climate of origin and the cultivation climate (i.e., greater genetic differentiation in plasticity) than structural traits. This implies that plasticity is trait specific and individuals from some plant populations will be more plastic and hence will be better able to persist under novel climates than others. Genetic controls over trait variability in plants in both structural (Münzbergová *et al*. 2017; Laitinen and Nikoloski 2019) and biochemical (Benomar *et al*. 2016; Kreyling *et al*. 2019) traits have been already reported, but to our knowledge, evidence for higher genetic differentiation in plasticity in biochemical traits than structural traits has not. This pattern may occur because biochemical traits are directly downstream of gene expression, able of quick and reversible responses while structural traits are one more step downstream than biochemical traits (i.e., structures are the product of biochemicals), are non-reversible, and require more time and energy to develop.

We also expected that plants originating from drier climates will show higher plasticity in traits. However, against our expectation, plants originating from climates with higher precipitation or a combination of higher temperature and precipitation tend to have higher plasticity in biochemical traits compared to structural traits. The possible reason for lower plasticity in plants originating from drier and colder climates could be that in such environments plants always tend to have high levels of photoprotective pigments (because of higher risk of photodamage) (Ramírez-Valiente *et al*. 2015). In optimal climatic conditions (i.e., with less abiotic filtering), there is no such requirement and plants may show more plastic responses. Studies also report that at warmer and wetter conditions competition becomes more prominent and higher plasticity provides an advantage to persist under high competition (Burns and Strauss 2012). Another possible reason could be attributed to trait convergence at colder and drier climates (Reich *et al*. 1997) due to which individuals originating from these climates have more similar trait values (i.e., lower plasticity). However, why it is more evident in biochemical traits is still an open question.

### Adaptiveness of plasticity depends upon climate (origin and cultivation) and trait type

Adaptive plasticity can facilitate or even accelerate the process of adaptive evolution (Ghalambor *et al*. 2007). We found evidence for adaptive as well as maladaptive plasticity in both types of traits. However, adaptive plasticity was more common in structural traits, while biochemical traits largely showed mal-adaptive plasticity (i.e., neutral or divergent response). These findings are in line with other earlier studies (van Kleunen and Fisher 2001; Ghalambor *et al*. 2007; Henn *et al*. 2018; Kreyling *et al*. 2019; Laitinen and Nikoloski 2019; Challis *et al*.2022; Brooker *et al*. 2022) suggesting that adaptive as well as mal-adaptive responses in plasticity might be quite common. Here, we report that the adaptiveness is also dependent on the type of plant function studied. A recent study (Arnold *et al*. 2022) also demonstrated that plastic responses to temperature in plants were highly dependent on the type of trait studied. Although biochemical traits are low-cost, reversible, and should show adaptive plasticity, we found the opposite. We speculate that when plants are growing outside climate of origin (i.e., non-optimal climates), they are under stress (Lichtenthaler 1998) which is causing divergence in biochemical traits. We also believe that greater genetic differentiation in plasticity in biochemical traits (Figure 4) could limit adaptive responses in biochemical traits.

Another important question is whether plants are showing adaptive plasticity in climates that are predicted for the future. In the context of climate change scenarios of the studied populations (i.e., in Norway), warming and increased precipitation are predicted (Hanssen-Bauer *et al*. 2017). By looking into our findings (related to adaptive plasticity in populations originating from cold and dry climates and cultivated in warm and wet climates), the species is likely to show adaptive plasticity in structural traits and will likely show mal-adaptive plasticity in photosynthesis related traits. Even if the climate change will be in the other direction (i.e., warmer and drier), the adaptive responses of populations will remain the same.

This combination of adaptive plasticity in structural and mal-adaptive plasticity in biochemical traits will likely lead to incomplete overall adaptive responses in plants (Ghalambor *et al*. 2007) and may have negative consequences on the persistence of the plant populations in the future (Lande 2009; Chevin *et al*. 2010). The ability of this species to adapt to the novel climate is also supported by Münzbergová et al. (2021), demonstrating that *Festuca rubra* (i.e., the same model species in the same study system as used here) is capable of real rapid evolution by selection of genotypes best adapted to novel conditions.

### Limitations of the study

Although this study reports important results on the variation in structural versus biochemical plant traits, it may be criticized because it uses only one replicate of each genotype in each cultivation climate, and therefore there is no replication at the genotype level. However, we did not aim to study the effects of the genotypes, but rather to control for their effects by having identical genotypes in each growth chamber. The lack of replication per genotype may especially affect the calculation of the ‘plasticity index’. However, 10 genotypes per population and, thus, 10 true replicates and independent plasticity estimates make the analyses robust.

## CONCLUSIONS

We conclude that the structural and biochemical traits of plants respond differentially to climate. Biochemical traits also represent an important but independent plant strategy linked to plant photosynthetic machinery i.e., *‘photoharvesting – photoprotection’* strategy dimension of functional variation. The responses of each trait type to climatic factors are complex and the many interactive effects of climate of origin and cultivation indicate that different populations will respond in a different way to novel climates. The traits showed adaptive as well as mal-adaptive plasticity. The adaptiveness also depends upon trait type and climate considered. Therefore, studying biochemical traits under multiple climates is important to better understand the effects of climate change on plants and predict the ability of plants to persist under climate change.

## ACKNOWLEDGEMENTS

We thank the POPEKOL discussion group for useful comments on the manuscript draft. The study was supported by the project GACR 19-00522S and institutional research projects RVO 67985939 and MSMT. We also thank Vigdis Vandvik and her team for granting access to the SEEDCLIM climate grid from where the initial material had been obtained. DT also acknowledges “European Union’s Horizon 2020 research and innovation programme under the Marie Sklodowska-Curie grant agreement No 101038052” for providing research fellowship.

## AUTHOR CONTRIBUTIONS

DT and ZM conceptualised the study. ZM supervised all the work including experiments and secured funding to carry out all the experimental work. DT analysed the data with extensive comments and suggestions from ZM. VH, RS, HS, DH and NW collected the experimental data. DT wrote the draft of the paper with extensive suggestions, inputs and editing from ZM. TD helped in some of the data analysis and provided comments during manuscript preparation. All the authors commented on and approved the final version of the manuscript.

