## SUPPLEMENTARY DATA for "Differential effect of climate on structural and biochemical plant traits"

**SUPPLEMENTARY DATA:** The variance partitioning model output (also summarized in Supplementary Table 3). Oprec and Otemp represent precipitation and temperature of climate of origin; Cprec and Ctemp represent precipitation and temperature of cultivation climate. Header represent the explanatory variable and the values in cells represent the variation explained by each variable

| Trait | Genotype | Otemp: Oprec:  Ctemp:  Cprec | Otemp:  Oprec:  Cprec | Otemp:  Oprec:  Ctemp | Oprec:  Ctemp:  Cprec | Otemp:  Ctemp:  Cprec | Oprec:  Cprec | Oprec:  Ctemp | Otemp:  Cprec |
| --- | --- | --- | --- | --- | --- | --- | --- | --- | --- |
| Aboveground biomass | 11 | 2.3 | 0 | 0 | 1.3 | 0 | 0 | 0 | 2.9 |
| Number of ramets | 24.85 | 4.72 | 0 | 0 | 0.15 | 0 | 1.81 | 0 | 2.87 |
| Rhizome weight | 13.78 | 0 | 4.54 | 0.06 | 0 | 0 | 0 | 0.93 | 0 |
| Root weight | 1.25 | 0.32 | 0 | 0.39 | 0.11 | 0 | 0 | 0.09 | 0 |
| Specific leaf area | 0.34 | 0.3 | 0 | 1.74 | 0 | 0 | 0.4 | 0 | 0 |
| Stomatal size | 32.68 | 0 | 0 | 1.85 | 0 | 5.69 | 0.58 | 0 | 0.87 |
| Stomatal density | 23.58 | 1.39 | 0 | 1.18 | 0 | 2.36 | 0 | 0 | 0 |
| Plant height | 11.02 | 0.63 | 0.56 | 0 | 0 | 0 | 0 | 0 | 0 |
| Neoxanthin | 2.1 | 8.62 | 1.55 | 5.56 | 3.56 | 0 | 17.75 | 3.48 | 12.93 |
| Violaxanthin | 3.48 | 20.82 | 7.38 | 0 | 0 | 0 | 11.5 | 3.31 | 1.04 |
| Antheraxanthin | 0.16 | 6.19 | 14.3 | 2.89 | 0 | 1.52 | 11.26 | 4.66 | 0 |
| Lutein | 0.4 | 10.21 | 4.56 | 3.39 | 4.59 | 0 | 17.31 | 0.64 | 0 |
| Zeaxanthin | 0.02 | 5.29 | 2.25 | 2.86 | 0 | 0.31 | 14.68 | 3.48 | 0 |
| �-carotene | 1.49 | 13.93 | 51.2 | 3.33 | 0 | 0.88 | 0 | 0 | 0 |
| Chlorophyll a | 0.86 | 7.42 | 0 | 0.52 | 3.23 | 0 | 26.65 | 5.78 | 5.35 |
| Chlorophyll b | 3.56 | 6.37 | 0.42 | 3.45 | 1.19 | 0 | 37.41 | 11.56 | 12.67 |
| Xanthophyll cycle de-epoxidation state (DEPS) | 0.24 | 6.79 | 0 | 1.59 | 0 | 0 | 15.25 | 3.89 | 0 |
| Phenolic compounds | 2.14 | 13.61 | 0.02 | 0 | 5.64 | 1.92 | 0 | 0 | 0 |
| Superoxide dismutase (SOD) | 39.63 | 0 | 0.92 | 0 | 0 | 0 | 3.38 | 0 | 2.28 |

| Trait | Otemp:Ctemp | Oprec | Otemp:Oprec | Otemp | Cprec | Ctemp | Ctemp:Cprec | Unexplained |
| --- | --- | --- | --- | --- | --- | --- | --- | --- |
| Aboveground biomass | 0 | 0 | 10 | 8.4 | 0 | 21 | 1.2 | 41 |
| Number of ramets | 0 | 6.96 | 0 | 0 | 9.08 | 0 | 0 | 49.57 |
| Rhizome weight | 0 | 0.89 | 0.44 | 0 | 44.43 | 0 | 0.39 | 34.54 |
| Root weight | 0 | 4.06 | 0.24 | 5.19 | 40.33 | 0 | 15.69 | 32.33 |
| Specific leaf area | 0.09 | 64.61 | 14.34 | 0 | 0 | 0.14 | 0 | 18.04 |
| Stomatal size | 0 | 0.78 | 0.11 | 0.45 | 0 | 0.61 | 0.05 | 56.34 |
| Stomatal density | 0 | 0 | 10.34 | 8.73 | 0 | 0 | 0.33 | 52.1 |
| Plant height | 0.11 | 1.16 | 0 | 0 | 1.83 | 47.08 | 16.7 | 20.91 |
| Neoxanthin | 0 | 0 | 18.21 | 0 | 15.24 | 0 | 1.22 | 9.77 |
| Violaxanthin | 0 | 0 | 0 | 0 | 35.82 | 2.7 | 0 | 13.95 |
| Antheraxanthin | 0 | 0 | 0 | 0 | 57.29 | 0 | 0 | 1.71 |
| Lutein | 0 | 0 | 7.45 | 0 | 44.48 | 0 | 1.34 | 5.63 |
| Zeaxanthin | 0 | 0 | 0 | 0 | 69.88 | 0 | 0 | 1.22 |
| �-carotene | 0 | 10.03 | 0 | 0 | 6.52 | 0.3 | 0 | 12.33 |
| Chlorophyll a | 1.16 | 0 | 0 | 0 | 42.71 | 0 | 0 | 6.34 |
| Chlorophyll b | 0 | 0 | 3.18 | 0 | 0 | 0 | 1.93 | 18.27 |
| Xanthophyll cycle de-epoxidation state (DEPS) | 0 | 0 | 0 | 0 | 70.51 | 0 | 0 | 1.74 |
| Phenolic compounds | 0 | 0 | 0 | 0 | 7.57 | 0 | 25.12 | 43.98 |
| Superoxide dismutase (SOD) | 0 | 0 | 1.79 | 0 | 12.02 | 0.49 | 1.82 | 37.68 |
