## SUPPLEMENTARY FIGURES AND TABLES for "Differential effect of climate on structural and biochemical plant traits"

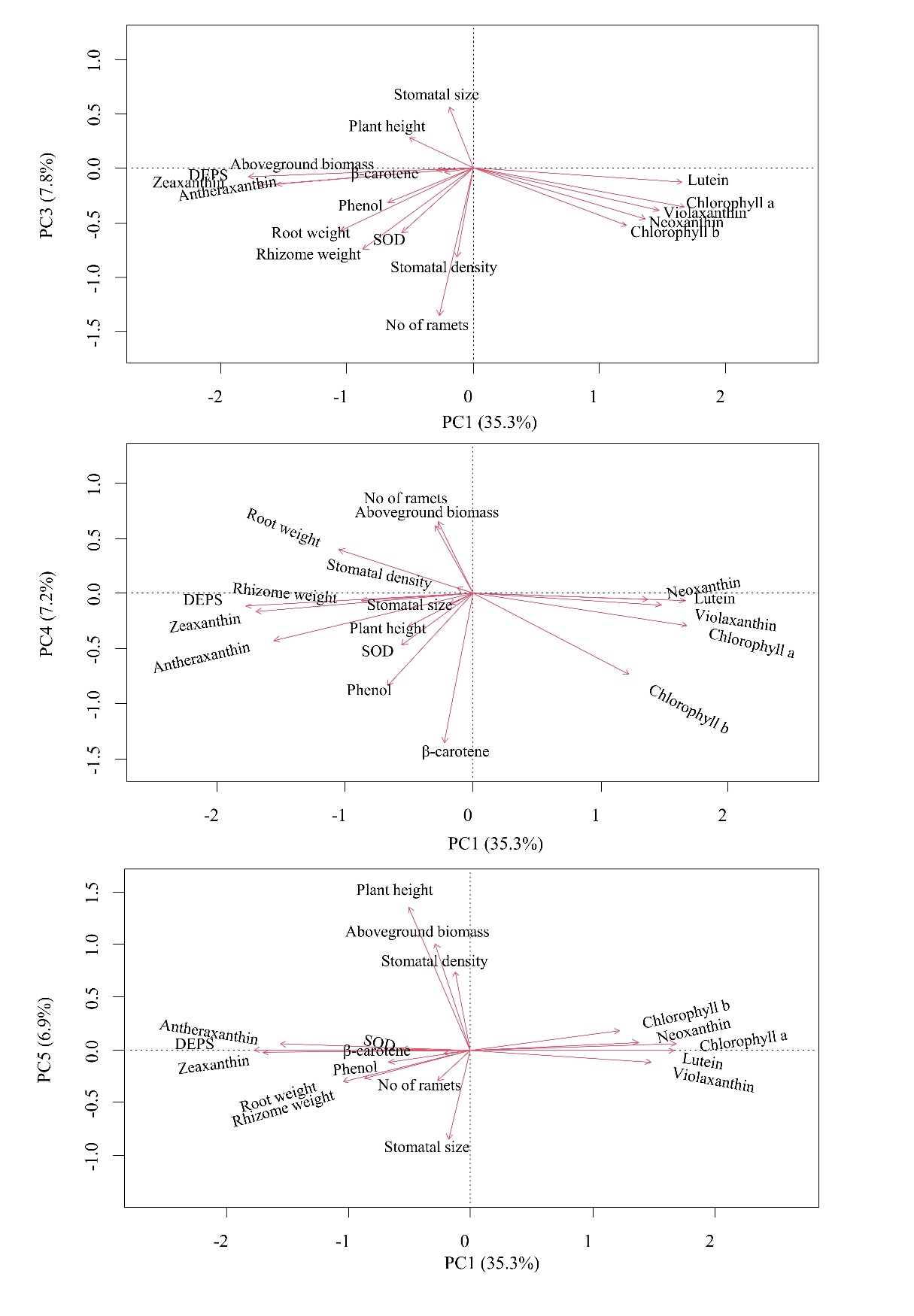

**Supplementary Figure 1:** Principal component (PC) patterns of studied functional traits. The axes 3 to 5 are plotted against axis 1. Plot of PC2 against PC1 is shown as Figure 1 in main text. DEPS = xanthophyll cycle de-epoxidation state; SOD = Superoxide dismutase.

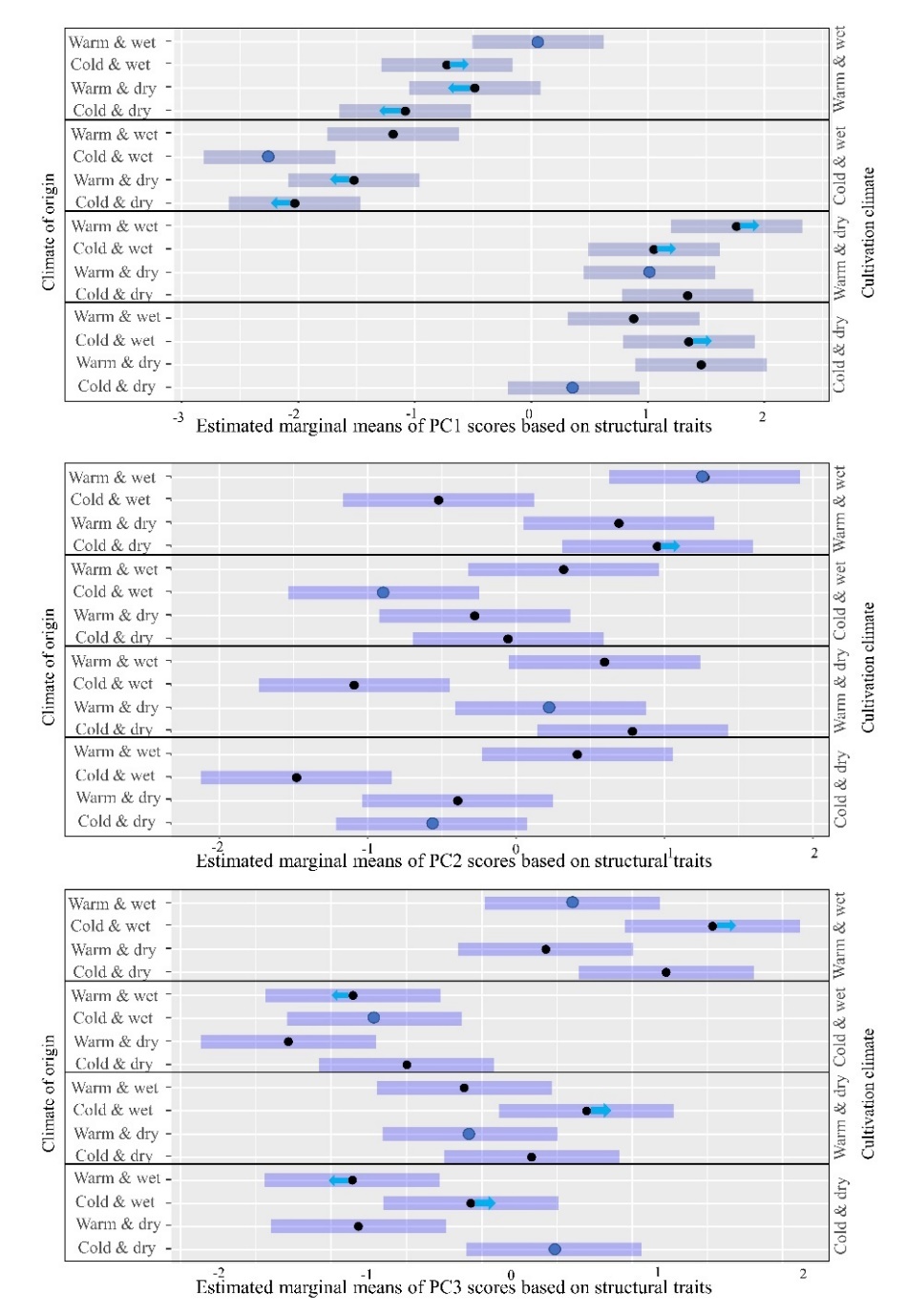

**Supplementary Figure 2:** Estimated marginal means of PC axis 1 to 3 (based on structural traits) for different populations cultivated in contrasting climates. The shaded areas are 95% confidence intervals. Means in large blue dots are for populations grown in climate of origin. The arrows on shaded areas represent direction of shift in traits (from climate of origin) along with convergence (blue arrows). This figure is based on analysis conducted using genotypes originating from the four most extreme localities.

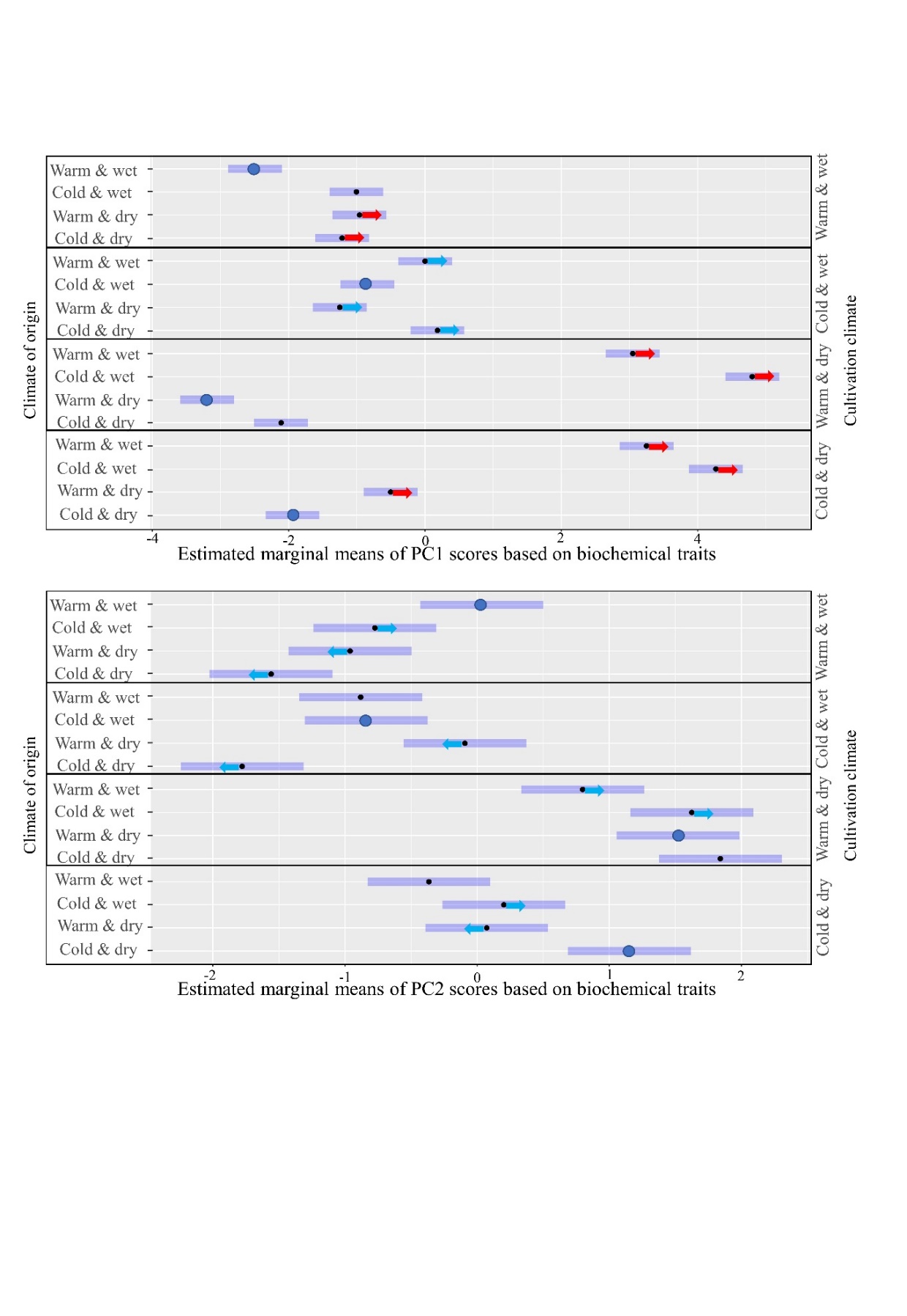

**Supplementary Figure 3:** Estimated marginal means of PCA axis 1 and 2 scores (based on biochemical traits) for different populations cultivated in contrasting climates. The shaded areas are 95% confidence intervals. Means in large blue dots are for populations grown in climate of origin. The arrows on shaded areas represent direction of shift in traits (from climate of origin) along with convergence (blue arrows) and divergence (red arrows). This figure is based on analysis conducted using genotypes originating from the four most extreme localities.

**Supplementary Table 1:** Trait loading and percentage of trait variation explained by the first five principal components (PCs). The upper half of the table is based on the data after imputation of missing values while the lower half is based on the dataset after omitting the samples with missing value for any of the trait. Loadings greater than 1 are in bold. Trait names in bold represent structural traits.

|  | **PC1** | **PC2** | **PC3** | **PC4** | **PC5** |
| --- | --- | --- | --- | --- | --- |
| ***Explained variation (%)*** | ***35.3*** | ***8.8*** | ***7.8*** | ***7.2*** | ***6.9*** |
| **Structural traits** | | | | | |
| Aboveground biomass | -0.34 | **1.4** | -0.01 | 0.72 | **1.18** |
| Plant height | -0.59 | 0.83 | 0.33 | -0.36 | **1.59** |
| Rhizome weight | **-1.02** | 0.1 | -0.88 | -0.07 | -0.32 |
| Root weight | **-1.23** | 0.92 | -0.67 | 0.47 | -0.36 |
| Stomatal density | -0.15 | **-1.27** | -0.96 | 0.06 | 0.87 |
| Stomatal size | -0.21 | **1.28** | 0.66 | -0.1 | **-1** |
| Number of ramets | -0.32 | 0.51 | **-1.58** | 0.77 | -0.35 |
| **Biochemical traits** | | | | | |
| Antheraxanthin | **-1.83** | 0.01 | -0.18 | -0.5 | 0.07 |
| β-carotene | -0.26 | 0.41 | -0.04 | **-1.59** | -0.04 |
| Chlorophyll *a* | **1.97** | 0.25 | -0.42 | -0.34 | 0.07 |
| Chlorophyll *b* | **1.43** | 0.11 | -0.62 | -0.87 | 0.22 |
| Xanthophyll cycle de-epoxidation state | **-2.09** | -0.17 | -0.09 | -0.14 | 0 |
| Phenolic compounds | -0.79 | 0.41 | -0.37 | -0.97 | -0.14 |
| Lutein | **1.95** | 0.17 | -0.15 | -0.08 | -0.01 |
| Neoxanthin | **1.61** | 0.44 | -0.54 | -0.06 | 0.08 |
| Superoxide dismutase | -0.66 | 0.05 | -0.7 | -0.55 | 0.01 |
| Violaxanthin | **1.73** | 0.41 | -0.45 | -0.12 | -0.14 |
| Zeaxanthin | **-2** | -0.1 | -0.18 | -0.19 | -0.03 |

|  | **PC1** | **PC2** | **PC3** | **PC4** | **PC5** |
| --- | --- | --- | --- | --- | --- |
| ***Explained variation (%)*** | ***35.6*** | ***8.8*** | ***7.9*** | ***7.4*** | ***6.9*** |
| **Structural traits** | | | | | |
| Aboveground biomass | -0.36 | **1.34** | -0.03 | 0.97 | -0.99 |
| Plant height | -0.57 | 0.78 | -0.59 | 0.10 | **-1.57** |
| Rhizome weight | -0.99 | 0.11 | 0.90 | -0.15 | 0.14 |
| Root weight | **-1.24** | 0.89 | 0.77 | 0.27 | 0.30 |
| Stomatal density | -0.11 | **-1.31** | 0.78 | 0.17 | -0.91 |
| Stomatal size | -0.22 | **1.32** | -0.50 | -0.27 | 0.97 |
| Number of ramets | -0.30 | 0.46 | **1.67** | 0.55 | 0.31 |
| **Biochemical traits** | | | | | |
| Antheraxanthin | **-1.81** | -0.02 | 0.11 | -0.48 | -0.20 |
| β-carotene | -0.28 | 0.33 | -0.14 | **-1.57** | -0.26 |
| Chlorophyll *a* | **1.95** | 0.25 | 0.37 | -0.31 | -0.20 |
| Chlorophyll *b* | **1.45** | 0.09 | 0.49 | -0.77 | -0.46 |
| Xanthophyll cycle de-epoxidation state | **-2.07** | -0.18 | 0.07 | -0.13 | -0.03 |
| Phenolic compounds | -0.78 | 0.45 | 0.28 | -1.06 | -0.18 |
| Lutein | **1.93** | 0.15 | 0.12 | -0.10 | -0.02 |
| Neoxanthin | **1.60** | 0.42 | 0.52 | -0.10 | -0.23 |
| Superoxide dismutase | -0.65 | 0.07 | 0.67 | -0.56 | -0.29 |
| Violaxanthin | **1.73** | 0.40 | 0.41 | -0.22 | 0.02 |
| Zeaxanthin | **-1.98** | -0.11 | 0.15 | -0.22 | -0.03 |

**Supplementary Table 2:** Redundancy analysis variable selection results showing change in R^2^ with addition of each variable to the model along with their significance. The explanatory variables that explain more than 1% of the variation (in bold) were retained in the final model.

| Explanatory Variables | R^2^ (cumulative) | F value | P value |
| --- | --- | --- | --- |
| **Cprec** | **0.21** | **119.37** | **0.001** |
| **+ Ctemp** | **0.25** | **21.85** | **0.001** |
| **+ Oprec** | **0.28** | **19.12** | **0.001** |
| **+ Oprec:Cprec** | **0.33** | **34.76** | **0.001** |
| **+ Ctemp:Cprec** | **0.35** | **15.94** | **0.001** |
| **+ Otemp** | **0.36** | **7.46** | **0.001** |
| + Otemp:Oprec | 0.37 | 5.15 | 0.001 |
| + Otemp:Ctemp | 0.37 | 3.11 | 0.001 |
| + Otemp:Cprec | 0.37 | 2.84 | 0.002 |
| + Oprec:Ctemp | 0.38 | 2.36 | 0.007 |
| + Oprec:Ctemp:Cprec | 0.38 | 2.68 | 0.005 |
| + Otemp:Ctemp:Cprec | 0.38 | 2.28 | 0.009 |
| + Otemp:Oprec:Ctemp | 0.38 | 2.02 | 0.015 |

**Supplementary Table 3:** Summary of variance partitioning analysis showing the variation explained in each trait by climate of origin, cultivation climate, their interaction and genotype. Significant effects are shown in bold. More detailed variation partitioning outputs are presented in Supplementary Data.

| Trait | Trait type | Origin | Cultivation | Origin × Cultivation | Genotype |
| --- | --- | --- | --- | --- | --- |
| **Structural traits** | | | | | |
| Aboveground biomass | ST | 18.42 | 22.22 | 6.52 | 11 |
| Number of ramets | ST | 6.98 | 9.1 | 9.57 | 24.85 |
| Rhizome weight | ST | 1.35 | 44.84 | 5.55 | 13.78 |
| Root weight | ST | 9.51 | 56.04 | 0.93 | 1.25 |
| Specific leaf area (SLA) | ST | 78.97 | 0.16 | 2.55 | 0.34 |
| Stomatal size | ST | 1.36 | 0.68 | 9.01 | 32.68 |
| Stomatal density | ST | 19.09 | 0.35 | 4.95 | 23.58 |
| Plant height | ST | 1.18 | 65.63 | 1.32 | 11.02 |
| **Biochemical traits** | | | | | |
| Neoxanthin | BC | 18.23 | 16.48 | 53.47 | 2.1 |
| Violaxanthin | BC | 0.02 | 38.54 | 44.07 | 3.48 |
| Antheraxanthin | BC | 0.02 | 57.31 | 40.84 | 0.16 |
| Lutein | BC | 7.47 | 45.84 | 40.72 | 0.4 |
| Zeaxanthin | BC | 0.02 | 69.9 | 28.89 | 0.02 |
| β-carotene | BC | 10.05 | 6.84 | 69.36 | 1.49 |
| Chlorophyll *a* | BC | 0.02 | 42.73 | 50.13 | 0.86 |
| Chlorophyll *b* | BC | 3.2 | 1.95 | 73.09 | 3.56 |
| Xanthophyll cycle de-epoxidation state (DEPS) | BC | 0.02 | 70.53 | 27.54 | 0.24 |
| Phenolic compounds | BC | 0.02 | 32.71 | 21.21 | 2.14 |
| Superoxide dismutase (SOD) | BC | 1.81 | 14.35 | 6.6 | 39.63 |

**Supplementary Table 4:** Trait loading and percentage of trait variation explained by the first three principal components (PCs) in structural and biochemical traits. The information in this table is based on plants originating from the four most extreme localities of SeedClim Grid.

| **Structural traits** | | | |
| --- | --- | --- | --- |
|  | **PC1** | **PC2** | **PC3** |
| ***Explained variation (%)*** | ***30.2*** | ***19.30*** | ***17.40*** |
| Aboveground biomass | 1.02 | -0.85 | 1.09 |
| Plant height | 0.79 | -0.91 | 1.45 |
| Rhizome weight | 1.18 | 1.11 | -0.38 |
| Root weight | 1.42 | -0.29 | -0.72 |
| Specific leaf area (SLA) | 1.69 | 0.36 | 0.26 |
| Stomatal density | 0.18 | 1.42 | 1.06 |
| Stomatal size | 0.68 | -1.28 | -0.86 |
| Number of ramets | 1.54 | 0.46 | -0.54 |
| **Biochemical traits** | | | |
|  | **PC1** | **PC2** | **PC3** |
| ***Explained variation (%)*** | ***55.70*** | ***15.40*** | ***9.20*** |
| Antheraxanthin | 1.62 | -0.67 | 0.51 |
| β-carotene | -1.67 | -0.50 | 0.12 |
| Chlorophyll *a* | -1.71 | -0.46 | 0.30 |
| Chlorophyll *b* | -1.08 | -0.93 | 1.03 |
| Xanthophyll cycle de-epoxidation state | 1.78 | -0.58 | 0.41 |
| Phenolic compounds | 0.03 | -1.13 | -1.34 |
| Lutein | -1.76 | 0.12 | 0.04 |
| Neoxanthin | -1.45 | -0.76 | 0.23 |
| Zeaxanthin | -1.54 | -0.42 | -0.22 |
| Violaxanthin | 1.68 | -0.81 | 0.37 |
| Superoxide dismutase | 0.51 | -1.28 | -0.47 |

**Supplementary Table 5:** Mixed effects model statistics showing the effect of climate of origin and cultivation climate on PC scores based structural and biochemical traits separately. Oprec and Otemp/Cprec and Ctemp represent precipitation and temperature of climate of origin/cultivation climate. Part A of the table presents variance and standard deviation of random effect, and estimate, standard error, and t value of fixed effects. Significant effects with F values and P values are shown in bold in Part B. This table is based on analysis based on genotypes originating from the four most extreme localities of SeedClim Grid.

**Part A**

| **Predictor** | **Structural traits** | | | | | | | | | **Biochemical traits** | | | | | |
| --- | --- | --- | --- | --- | --- | --- | --- | --- | --- | --- | --- | --- | --- | --- | --- |
|  | **PC1** | | | **PC2** | | | **PC3** | | | **PC1** | | | **PC2** | | |
| ***Random Effect*** | ***Var.*** | ***SD*** |  | ***Var.*** | ***SD*** |  | ***Var.*** | ***SD*** |  | ***Var.*** | ***SD*** |  | ***Var.*** | ***SD*** |  |
| Genotype (Intercept) | 0.12 | 0.35 |  | 0.13 | 0.36 |  | 0.39 | 0.63 |  | 0.13 | 0.36 |  | 0.21 | 0.46 |  |
| Residual | 0.69 | 0.83 |  | 0.93 | 0.96 |  | 0.46 | 0.68 |  | 0.27 | 0.51 |  | 0.34 | 0.58 |  |
| ***Fixed effects*** | ***Est.*** | ***SE*** | ***t value*** | ***Est.*** | ***SE*** | ***t value*** | ***Est.*** | ***SE*** | ***t value*** | ***Est.*** | ***SE*** | ***t value*** | ***Est.*** | ***SE*** | ***t value*** |
| Intercept | 0.36 | 0.28 | 1.28 | -0.57 | 0.33 | -1.75 | 0.48 | 0.29 | 1.64 | -1.94 | 0.20 | -9.78 | 1.15 | 0.23 | 4.92 |
| Otemp | 1.09 | 0.40 | 2.72 | 0.18 | 0.46 | 0.39 | -1.23 | 0.41 | -3.14 | 1.44 | 0.28 | 5.12 | -1.08 | 0.33 | -3.26 |
| Oprec | 0.99 | 0.40 | 2.46 | -0.91 | 0.46 | -1.98 | -0.55 | 0.41 | -1.33 | 6.21 | 0.28 | 22.09 | -0.95 | 0.33 | -2.87 |
| Ctemp | 0.98 | 0.37 | 2.64 | 1.36 | 0.43 | 3.15 | -0.15 | 0.30 | -0.49 | -0.17 | 0.23 | -0.73 | 0.69 | 0.26 | 2.64 |
| Cprec | -2.39 | 0.37 | -6.45 | 0.52 | 0.43 | 1.20 | -0.98 | 0.30 | -3.21 | 2.13 | 0.23 | 9.24 | -2.93 | 0.26 | -11.24 |
| Otemp:Oprec | -1.57 | 0.57 | -2.75 | 1.72 | 0.65 | 2.64 | 0.51 | 0.58 | 0.88 | -2.46 | 0.40 | -6.18 | 0.51 | 0.47 | 1.10 |
| Otemp:Ctemp | -1.42 | 0.53 | -2.71 | -0.73 | 0.61 | -1.20 | 0.89 | 0.43 | 2.07 | -2.52 | 0.33 | -7.75 | 0.76 | 0.37 | 2.06 |
| Otemp:Cprec | -0.59 | 0.53 | -1.12 | -0.40 | 0.61 | -0.66 | 0.51 | 0.43 | 1.19 | -2.87 | 0.33 | -8.82 | 2.76 | 0.37 | 7.50 |
| Oprec:Ctemp | -1.28 | 0.53 | -2.43 | -0.97 | 0.61 | -1.59 | 0.92 | 0.43 | 2.13 | 0.70 | 0.33 | 2.15 | 0.73 | 0.37 | 1.99 |
| Oprec:Cprec | -1.20 | 0.53 | -2.29 | 0.07 | 0.61 | 0.12 | 0.34 | 0.43 | 0.78 | -7.24 | 0.33 | -22.24 | 1.89 | 0.37 | 5.13 |
| Ctemp:Cprec | -0.03 | 0.53 | -0.06 | -0.35 | 0.61 | -0.57 | 1.87 | 0.43 | 4.35 | -1.23 | 0.33 | -3.78 | -0.47 | 0.37 | -1.27 |
| Otemp:Oprec:Ctemp | 2.60 | 0.74 | 3.51 | 0.53 | 0.86 | 0.61 | -0.92 | 0.61 | -1.51 | 1.79 | 0.46 | 3.89 | -1.02 | 0.52 | -1.96 |
| Otemp:Oprec:Cprec | 2.12 | 0.74 | 2.86 | -0.28 | 0.86 | -0.32 | 0.13 | 0.61 | 0.22 | 4.74 | 0.46 | 10.29 | -2.24 | 0.52 | -4.30 |
| Otemp:Ctemp:Cprec | 1.51 | 0.74 | 2.03 | 0.69 | 0.86 | 0.81 | -0.90 | 0.61 | -1.49 | 4.21 | 0.46 | 9.15 | -1.85 | 0.52 | -3.55 |
| Oprec:Ctemp:Cprec | 1.85 | 0.74 | 2.49 | 0.33 | 0.86 | 0.38 | -0.39 | 0.61 | -0.65 | 0.54 | 0.46 | 1.17 | -0.89 | 0.52 | -1.70 |
| Otemp:Oprec:Ctemp:Cprec | -2.97 | 1.05 | -2.83 | 0.09 | 1.22 | 0.08 | 0.14 | 0.86 | 0.16 | -5.82 | 0.65 | -8.93 | 2.96 | 0.74 | 4.01 |

**Part B**

| Predictor | **Structural traits** | | | | | | **Biochemical traits** | | | |
| --- | --- | --- | --- | --- | --- | --- | --- | --- | --- | --- |
|  | PC1 | | PC2 | | PC3 | | PC1 | | PC2 | |
|  | **F value** | **P value** | **F value** | **P value** | **F value** | **P value** | **F value** | **P value** | **F value** | **P value** |
| Otemp | 8.30 | **0.007** | 14.15 | **<0.001** | 10.98 | **0.002** | 628 | **<0.001** | 91.6 | **<0.001** |
| Oprec | 1.90 | 0.177 | 3.27 | 0.079 | 0.06 | 0.812 | 1.6 | 0.207 | 0.3 | 0.614 |
| Ctemp | 31.24 | **<0.001** | 23.1 | **<0.001** | 115.3 | **<0.001** | 628 | **<0.001** | 91.6 | **<0.001** |
| Cprec | 308.2 | **<0.001** | 6.07 | **0.015** | 1.03 | 0.313 | 25.5 | **<0.001** | 13.6 | **<0.001** |
| Otemp:Oprec | 0.02 | 0.881 | 23.9 | **<0.001** | 0.12 | 0.735 | 0.6 | 0.425 | 0.4 | 0.542 |
| Otemp:Ctemp | 0.17 | 0.683 | 0.10 | 0.749 | 0.01 | 0.942 | 628.1 | **<0.001** | 91.6 | **<0.001** |
| Otemp:Cprec | 3.43 | 0.067 | 0.31 | 0.577 | 0.56 | 0.457 | 26.3 | **<0.001** | 13.6 | **<0.001** |
| Oprec:Ctemp | 0.63 | 0.431 | 2.88 | 0.093 | 1.89 | 0.172 | 7.4 | **0.008** | 5.1 | **0.026** |
| Oprec:Cprec | 0.02 | 0.884 | 0.15 | 0.695 | 1.26 | 0.265 | 24.5 | **<0.001** | 5.7 | **0.019** |
| Ctemp:Cprec | 11.97 | **<0.001** | 0.38 | 0.540 | 34.10 | **<0.001** | 30.7 | **<0.001** | 5.9 | **0.017** |
| Otemp:Oprec:Ctemp | 4.54 | 0.035 | 0.88 | 0.351 | 3.88 | 0.052 | 0 | 0.939 | 2.4 | 0.126 |
| Otemp:Oprec:Cprec | 1.46 | 0.230 | 0.14 | 0.704 | 0.22 | 0.641 | 19.5 | **<0.001** | 4.7 | **0.032** |
| Otemp:Ctemp:Cprec | 0.00 | 0.963 | 1.48 | 0.227 | 3.76 | 0.055 | 53.7 | **<0.001** | 14.3 | **<0.001** |
| Oprec:Ctemp:Cprec | 0.49 | 0.487 | 0.38 | 0.539 | 0.56 | 0.454 | 10.4 | **0.002** | 9.9 | **0.002** |
| Otemp:Oprec:Ctemp:Cprec | 8.01 | **0.006** | 0.01 | 0.940 | 0.03 | 0.872 | 79.8 | **<0.001** | 16.1 | **<0.001** |

**Supplementary Table 6:** Summary of the redundancy analysis testing the dependence of traits on climate of origin and cultivation. Part A of the table represents biplot scores while part B represents the model statistics.

**Part A**

| ***Biplot scores for different traits*** | | |
| --- | --- | --- |
|  | CAP1 | CAP1 |
| **Structural traits** | | |
| Aboveground biomass | 0.26 | -0.65 |
| Plant height | 0.58 | -1.41 |
| Rhizome weight | 0.94 | -0.06 |
| Root weight | 1.24 | 0.23 |
| Stomatal density | 0.14 | 0.09 |
| Stomatal size | 0.18 | -0.24 |
| Number of ramets | 0.39 | -0.10 |
| **Biochemical traits** | | |
| Antheraxanthin | 1.57 | -0.02 |
| β-carotene | 0.22 | -0.59 |
| Chlorophyll *a* | -1.70 | -0.14 |
| Chlorophyll *b* | -1.15 | -0.29 |
| Xanthophyll cycle de-epoxidation state | 2.0 | 0.15 |
| Phenolic compounds | 0.82 | -0.25 |
| Lutein | -1.63 | -0.11 |
| Neoxanthin | -1.12 | -0.09 |
| Zeaxanthin | -1.35 | 0.16 |
| Violaxanthin | 2.00 | 0.20 |
| Superoxide dismutase | 0.64 | -0.18 |
| ***Biplot scores for constraining variables*** | | |
|  | CAP1 | CAP2 |
| Tprec | -0.86 | 0.09 |
| Ttemp | 0.20 | -0.85 |
| Oprec | 0.26 | 0.35 |
| Otemp | -0.01 | -0.08 |
| Tprec:Oprec | -0.63 | 0.15 |
| Tprec:Ttemp | -0.55 | -0.50 |

**Part B**

| Explanatory Variables | Variance | F value | P value |
| --- | --- | --- | --- |
| Cprec | 3.72 | 140.8 | 0.001 |
| Ctemp | 0.71 | 26.72 | 0.001 |
| Oprec | 0.59 | 22.14 | 0.001 |
| Otemp | 0.18 | 6.63 | 0.001 |
| Cprec:Oprec | 1.01 | 38.03 | 0.001 |
| Cprec:Ctemp | 0.37 | 13.84 | 0.001 |

**Supplementary Table 7:** Summary of the redundancy analysis testing the dependence of traits on climate differences. Part A of the table represents biplot scores while part B represents the model statistics.

**Part A**

| ***Biplot scores for different traits*** | | |
| --- | --- | --- |
|  | CAP1 | CAP2 |
| **Structural traits** | | |
| Aboveground biomass | 0.18 | 0.46 |
| Plant height | 0.44 | 1.04 |
| Rhizome weight | 0.89 | -0.10 |
| Root weight | 1.10 | -0.59 |
| Stomatal density | 0.34 | 0.22 |
| Stomatal size | 0.1 | -0.08 |
| Number of ramets | 0.38 | -0.13 |
| **Biochemical traits** | | |
| Antheraxanthin | 1.36 | 0.001 |
| β-carotene | -0.03 | 0.27 |
| Chlorophyll *a* | -1.36 | 0.04 |
| Chlorophyll *b* | -0.94 | 0.08 |
| Xanthophyll cycle de-epoxidation state | 1.75 | 0.05 |
| Phenolic compounds | 0.94 | 0.10 |
| Lutein | -1.34 | 0.09 |
| Neoxanthin | -0.77 | -0.06 |
| Zeaxanthin | -1.0 | -0.13 |
| Violaxanthin | 1.81 | 0.06 |
| Superoxide dismutase | 0.72 | -0.002 |
| ***Biplot scores for constraining variables*** | | |
|  | CAP1 | CAP2 |
| ChangeT | 0.18 | 0.96 |
| ChangeP | -0.98 | 0.11 |
| ChangeT:ChangeP | -0.13 | 0.23 |

**Part B**

|  | Variance | F value | P value |
| --- | --- | --- | --- |
| ChangeP | 3.47 | 109.4 | 0.001 |
| ChangeT | 0.36 | 14.6 | 0.001 |
| ChangeP:ChangeT | 0.25 | 7.94 | 0.001 |
