## SUPPLEMENTARY METHODS for "Differential effect of climate on structural and biochemical plant traits"

During the experiment, in mid-June, the biomass of the plants were cut at 3 cm to simulate biomass removal during regular management in the field sites. This biomass was used for trait measurements as described below. The number of ramets was counted and the length (cm) of the longest ramet (hereafter referred to as plant height) was measured at the end of August 2015 and the plants were subsequently harvested, sorted into aboveground parts, roots and rhizomes, dried to a constant mass at 60 °C and weighed.

The remaining traits were measured during biomass removal in mid-June 2015. Stomatal density and stomatal size (length) were measured on epidermal imprints generated using nail polish (Gitz & Baker, 2009). The impressions were mounted on microscope slides with transparent adhesive tape and stomata were counted from three different regions on a leaf (each 500 × 500μm). From each of these three areas, three stomata were randomly chosen, and their length was measured, resulting in nine measured stomata per sample. SLA was estimated as a ratio of leaf area (mm^2^) to dry mass (mg). Leaf area was calculated on scanned folded leaves (as *Festuca* leaves are naturally folded and are often hard to unfold) multiplied by two to acquire the real area of the leaves.

The analyses of chlorophylls and carotenoids followed the protocol described in (Münzbergová and Haisel 2019). The freshly sampled leaves (collected from the plants in mid-June 2015) were frozen in liquid nitrogen, lyophilized for 24 h and stored in a freezer (-80 °C) until the analyses. The contents of chlorophyll (chlorophyll a and chlorophyll b) and carotenoids (b-carotene, lutein, neoxanthin, violaxanthin, antheraxanthin, and zeaxanthin) in all the samples were analyzed by High-performance liquid chromatography (HPLC; ECOM, Prague, Czech Republic) using a reversed-phase column (Watrex Nucleosil 120-5-C18, 5 mm particle size, 125 x 4 mm, ECOM, Prague, Czech Republic). Prior to HPLC injection, pigments were extracted from the leaves with acetone. In HPLC, elution was carried out for 25 min using a gradient solvent system acetonitrile/methanol/water (80:12:6) followed by methanol:ethylacetate (9:1), the gradient was run at the time of 2–5 min. The flow rate was 1 ml min^-1^ and the detection wavelength was 445 nm. Each sample was analyzed twice, and the results were averaged. The contents of all the pigments were expressed as mg/g of dry weight. The de-epoxidation state (DEPS) was derived from xanthophyll pigments following (Pospíšilová et al. 2000).

Total content of phenolic compounds was quantified spectrophotometrically from 80% methanolic extract of dried leaves using modified Folin-Ciocalteu method (Folin and Ciocalteu 1927). The absorbance was measured in 1 cm cuvette at 765 nm. The total phenolic compound concentration in the analyzed extract was expressed in gallic acid equivalents.

SOD was extracted from frozen leaves homogenized with mortar and pestle in 2.5 ml buffer (0.1 M Tris-HCl, 1 mM dithiothreitol, 1 mM Na2EDTA, 1% Triton X-100, 5 mM ascorbic acid, pH 7.8) and a small amount of PVP (polyvinylpyrrolidone). Samples were incubated on ice in the dark for 30 min and centrifuged (16,000 g, 10 min, 4 °C). Pellets were discarded, and supernatants were divided into Eppendorf tubes, frozen in liquid nitrogen, and stored at -70 °C (Lubovska et al., 2014). Later, supernatants were used for gel electrophoresis and SOD was stained on the gel (Fridovich 1986). These stained gels were scanned and densitograms were created and analyzed in the ImageJ program (Schneider et al. 2012). The relative activity of total SOD was estimated as the sum of intensities of bands expressed as peak areas.
